# Regulation of Aortic Morphogenesis and VE-Cadherin Dynamics by VEGF

**DOI:** 10.1101/2022.06.09.495466

**Authors:** Julian Jadon, Ronit Yelin, Alaa A. Arraf, Manar Abboud Asleh, Mira Zaher, Thomas M. Schultheiss

## Abstract

In amniote vertebrates, the definitive dorsal aorta is formed by the side-to-side fusion of two primordial aortic endothelial tubes. Formation of the definitive dorsal aorta requires extensive cellular migrations and rearrangements of the primordial tubes in order to generate a single fused vessel located at the embryonic ventral midline. This study examines the role of VEGF signaling in the generation of the definitive dorsal aorta. Through gain- and loss-of-function studies in vivo in the chick embryo, we document a requirement for VEGF signaling in growth and migration of the paired primordia, and identify a developmental window in which aorta development is particularly sensitive to VEGF signaling. We also find that VEGF signaling regulates the intracellular distribution between membrane and cytoplasm of the cell-cell adhesion molecule VE-Cadherin in aortic endothelial cells in vivo, suggesting a possible mechanism for how VEGF regulates aortic endothelial cell dynamics during the fusion process.

## Introduction

The dorsal aorta is the largest blood vessel in vertebrates and an equivalent structure is also found in many invertebrates. In vertebrates, the dorsal aorta, like other blood vessels, is derived from a collection of mesodermal endothelial precursors (angioblasts) located in the splanchnic layer of the lateral plate mesoderm (Drake and Fleming, 2000; Risau and Flamme, 1995; Sato, 2013). Angioblasts assemble into primitive vascular networks that remodel, eventually forming the dorsal aortae (Hirakow and Hiruma, 1981; Sato, 2013). The aorta initially appears as two endothelial tubes ventral to the intermediate mesoderm on both sides of the embryo. During the course of development, the two aortae move medially and fuse, forming the descending dorsal aorta located in the midline of the embryo ventral to the notochord (Fig. 1)(Sato, 2013). This fusion occurs in the main trunk region of the body. Posterior to the point where the large vitelline arteries exit laterally towards the yolk sac, the aortae remain unfused and extend into the more posterior regions of the body as two separate vessels.

**Figure 1.**
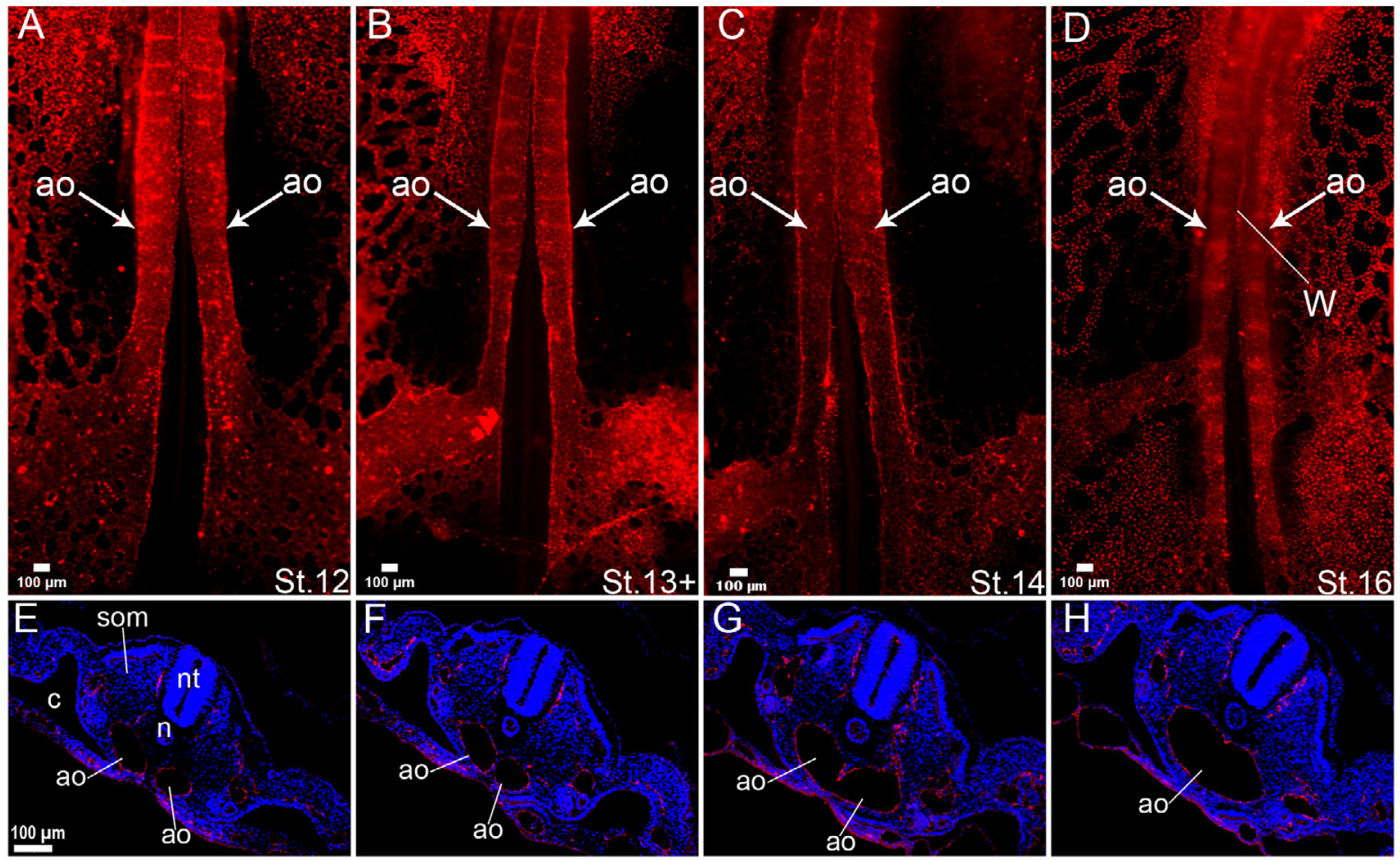
Overview of Aortic Fusion. (A-D) Whole-mount views of embryos injected with DiI-LdL into the heart to label all endothelial cells. Note that the aortae become increasingly close to each other and that fusion occurs from anterior (top) to posterior (bottom). “w” indicates the shared wall where the aortae have made contact with each other. (E-H) Cross sections of embryos at successive stages of midline convergence and fusion, stained with DiI-LdL (red) and with Dapi (blue) to mark nuclei. In E the aortae are still separated from each other, in F they have made contact, in G the common wall between the aorta is partially broken down, and in H the common wall is completely absent. Ao, aorta; c, coelomic cavity; n, notochord; nt, neural tube; som, somite.

The processes by which the dorsal aortae are created, move medially, and fuse into a single vessel have received some attention in the scientific literature but are still far from being fully understood (Meadows et al., 2012; Reese et al., 2004). BMP signaling is required for medial migration of the aortae; BMP inhibitors secreted by midline structures prevent medial aorta migration and are down-regulated during development to allow migration to proceed (Bressan et al., 2009; Garriock et al., 2010). However it is not yet clear how BMP signaling promotes aorta migration at the cellular level. In Xenopus, VEGF secreted from the midline hypochord is required for medial aorta migration (Cleaver and Krieg, 1998); however amniote vertebrates do not have a hypochord and it is not clear whether an equivalent midline source of VEGF signaling exists.

The current study focuses on the role of VEGF signaling during formation and morphogenesis of the amniote aorta, using the chick embryo as a model system. Vascular Endothelial Growth Factor (VEGF) is considered to be a master harmonizer of the initiation and maintenance of blood vessels (Gerhardt et al., 2003; Plouët et al., 1989). Secreted VEGF binds to two main tyrosine kinase receptors, VEGFr1 (Flt1) and VEGFr2 (KDR/Flk-1), which are both specifically expressed in endothelial cells (Neufeld et al., 1999). A naturally occurring truncated, secreted form of Flt1 (sFlt1) can act as a VEGF inhibitor (Maynard et al., 2003). VEGF binding to VEGFr2 on endothelial cells promotes endothelial cell division, and in addition promotes chemotaxis towards the VEGF source. VEGF signaling through VEGFr2 also initiates actin polymerization (Lamalice et al., 2007), generating filopodia extensions that enlarge cell surface area and expose cells to more VEGF ligand. Knockout mice for VEGF or either of its receptors are embryonic lethal, exhibiting severe loss of vasculogenesis (Ferrara et al., 1996; Fong et al., 1995; Shalaby et al., 1995). VEGF exhibits a strong dose response, as heterozygote deletions of VEGF produce strong vascular disruption and loss of vascular structures (Carmeliet et al., 1996; Ferrara et al., 1996).

Because of the strong early phenotype of disruption of VEGF signaling, it has been difficult to gather information on later roles of the VEGF pathway during embryonic vessel formation. In the current study we address the role of VEGF signaling on aorta morphogenesis in the chick embryo, where targeted in vivo manipulation at later embryonic stages is feasible.

Blood begins to circulate in the chick embryo at about HH Stage 12, prior to aortic fusion. This implies that mechanisms must exist to maintain endothelial wall integrity despite the extensive morphogenesis that the aortae undergo during this period. VE-Cadherin (VE-Cad), an endothelial cell-specific cadherin, is a prominent component of endothelial cell-cell junctions (Vestweber, 2008). VE-Cad is linked through catenin proteins and vinculin to the actin cytoskeleton, and thus VE-Cad mediates a strong link between the cytoskeleton of adjacent endothelial cells (Carmeliet et al., 1999). Ve-cad is also found to interact with other molecules to promote contact inhibition of growth (Dejana et al., 2009). Expression of VEGF in Human Umbilical Vein Endothelial Cells (HUVECS) induced the endocytosis of VE-Cad via clathrin-dependent endocytosis (Gavard and Gutkind, 2006; Hebda et al., 2013), suggesting a possible link between VEGF signaling and cell-cell interactions mediated by VE-Cadherin. One goal of the current study was to investigate the effects of VEGF signaling on VE-Cad dynamics in an in vivo context during aorta morphogenesis.

The results reported here point to strong effects of VEGF signaling on multiple stages of aortic morphogenesis, including vessel formation and maintenance, migration, and fusion, and reveal a VEGF-induced intracellular redistribution of VE-Cadherin that may mediate some of these effects.

## Results

Between days 2 and 3 of development, the bilateral dorsal aortae, which are initially located ventral to the intermediate mesoderm (IM), converge medially towards the embryonic midline, where they fuse to form a single dorsal aorta vessel that in located between the notochord and the endoderm (Fig. 1). Fig. 1A-D show of different stages of embryos in which all blood vessels were labeled by the fluorescent tracer DiI-LdL, which was injected into the beating heart. DiI-LdL is endocytosed specifically by endothelial cells, which express scavenger LdL receptors (Adachi et al., 1997). Fig. 1E-H show representative cross sections of the same embryo at different axial levels showing stages of the fusion process. Fusion occurs in an anterior-to-posterior wave, accompanied by break down of the common wall that separates the two aortae (compare arrows in Fig. 1F and Fig. 1G,H).

In order to investigate potential roles for VEGF signaling in aorta dynamics, we first used in situ hybridization to examine the expression of mRNA’s encoding for VEGF and its ligand VEGFR2/Flk1. As seen in Fig. 2D-F, Flk1 is expressed in all endothelial cells and is specific for endothelial cells. VEGF ligand is expressed in the endoderm, primarily in the medial regions, in the splanchnopleuric mesoderm, and in the ventro-medial regions of somites. At early stages of aorta medial convergence (Fig. 2A,B), the aortic endothelium is surrounded by VEGF-expressing tissues on its medial, lateral, and dorsal sides, although not on its ventral side, where it is in contact with endoderm. Subsequent medial movements bring the ventral wall of the aorta into contact also with VEGF-expressing endoderm (Fig. 2C).

**Figure 2.**
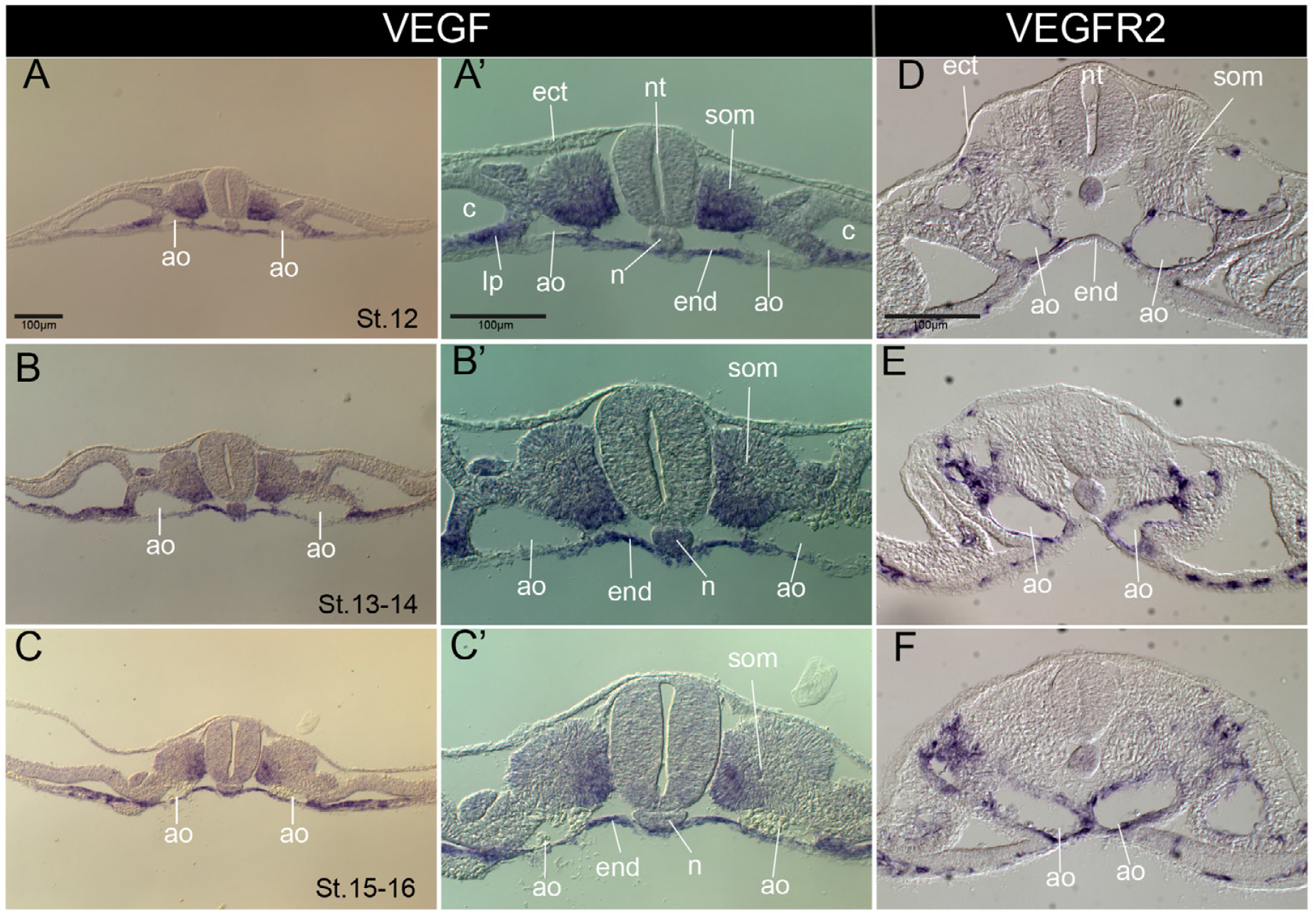
Expression of VEGF and VEGFR2. In situ hybridization for mRNA of VEGF (A-C) and VEGFR2 (D-F). A’-C’ are magnifications of A-C, respectively. Ao, aorta; c, coelomic cavity; ect, ectoderm; end, endoderm; n, notochord; nt, neural tube; som, somite.

In order to investigate roles for VEGF signaling in aorta dynamics, we made use of the secreted, truncated VEGF receptor sFlt1, a naturally occurring truncated form of the receptor Flt1, which binds VEGF extracellularly and prevents it from binding to its native receptors, thus lowering available VEGF ligand levels (Maynard et al., 2003). A plasmid co-expressing sFlt1 and GFP was electroporated into the endoderm of HH St. 12 embryos, from which it was presumably secreted into surrounding tissues. As seen in Fig. 3B,B’,J) sFlt1-electroporated embryos exhibited delayed aortic fusion (compare with control embryos electroporated with eGFP alone, Fig. 3A,A’,I). In many cases, sFlt1 caused degeneration of the aortae such that they became much smaller and without lumens (Fig. 3F, compare with control Fig. 3E), while in other cases the aortae appeared smaller but intact (Fig. 3I,J). These observations suggest that the delay/interference with aortic fusion may be caused, at least in part, by defects in aortic growth, and that aortic growth and medial movement are linked.

**Figure 3.**
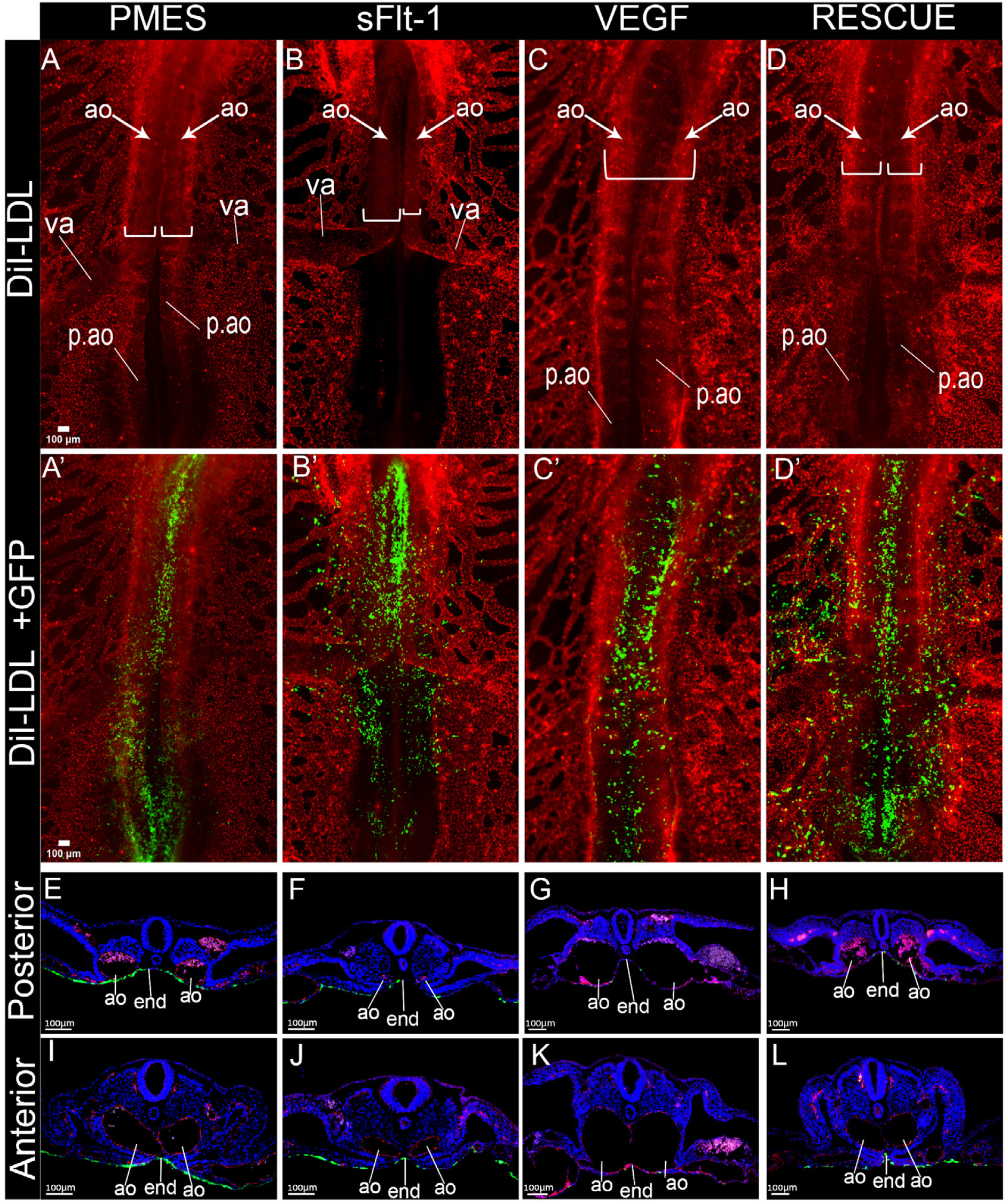
Lowering of VEGF concentration prevents aorta development and fusion. The endoderm was electroporated with control plasmid pMES (A,A’,E,I), sFlt1 (B,B’,F,J); VEGF (C,C’,G,K), and sFlt1 plus VEGF (D,D’,H,L). A-D and A’-D’ are whole mount views stained for DiI-LdL to mark blood vessels (A-D and A’-D’) and Gfp to mark electroporated cells (A’-D’). E-L are cross sections taken from posterior (E-H) or anterior (I-L) regions and stained for VE-Cadherin (red) to mark blood vessels (red) and Gfp to mark electroporated cells. There is also autofluorescence of blood cells in the red channel. Brackets in A-D indicate width of aortae. Note absence of aorta posterior to the vitelline artery in B. Ao, aorta; end, endoderm; p.ao, aorta posterior to 22^nd^ somite; va, vitelline artery.

In complementary experiments, plasmid encoding a processed form of VEGF (VEGF165) was electroporated into the endoderm. VEGF caused a marked growth in the aorta relative to other parts of the embryo (Fig. 3C,G,K), and appeared to cause a premature breakdown of the common wall between the two aortae (Fig. 3K). Co-electroporation of sFlt1 and VEGF restored a relatively normal phenotype (Fig. 3D,H,L), pointing to the specificity of both the sFlt1 and VEGF effects.

The effects of sFlt1 and VEGF on aorta convergence and fusion prompted us to look for other effects of manipulating VEGF levels on aorta dynamics. In the normal arrangement of large embryonic vessels, the aorta makes a connection at the axial level of the 22^nd^ somite to the vitelline artery (VA), the major link between the intraembryonic and extraembryonic circulations (Fig. 4A). The aorta also extends posteriorly beyond the 22^nd^ somite into the posterior regions of the embryo, where the aortae remain unfused (Fig. 4A). When embryos are electroporated with sFlt1, extension of the aorta posteriorly beyond the junction with the VA is inhibited (Fig. 4B). In most cases, when embryos are stained with antibodies to the endothelial marker VE-Cadherin (VE-Cad), endothelial structures can be found in the normal location of the posterior aorta (Fig. 4C), but these are not connected to the main aortic vessel and thus are not labeled by injection of DiI-LdL into the circulating blood (Fig. 4B-B’’).

**Figure 4.**
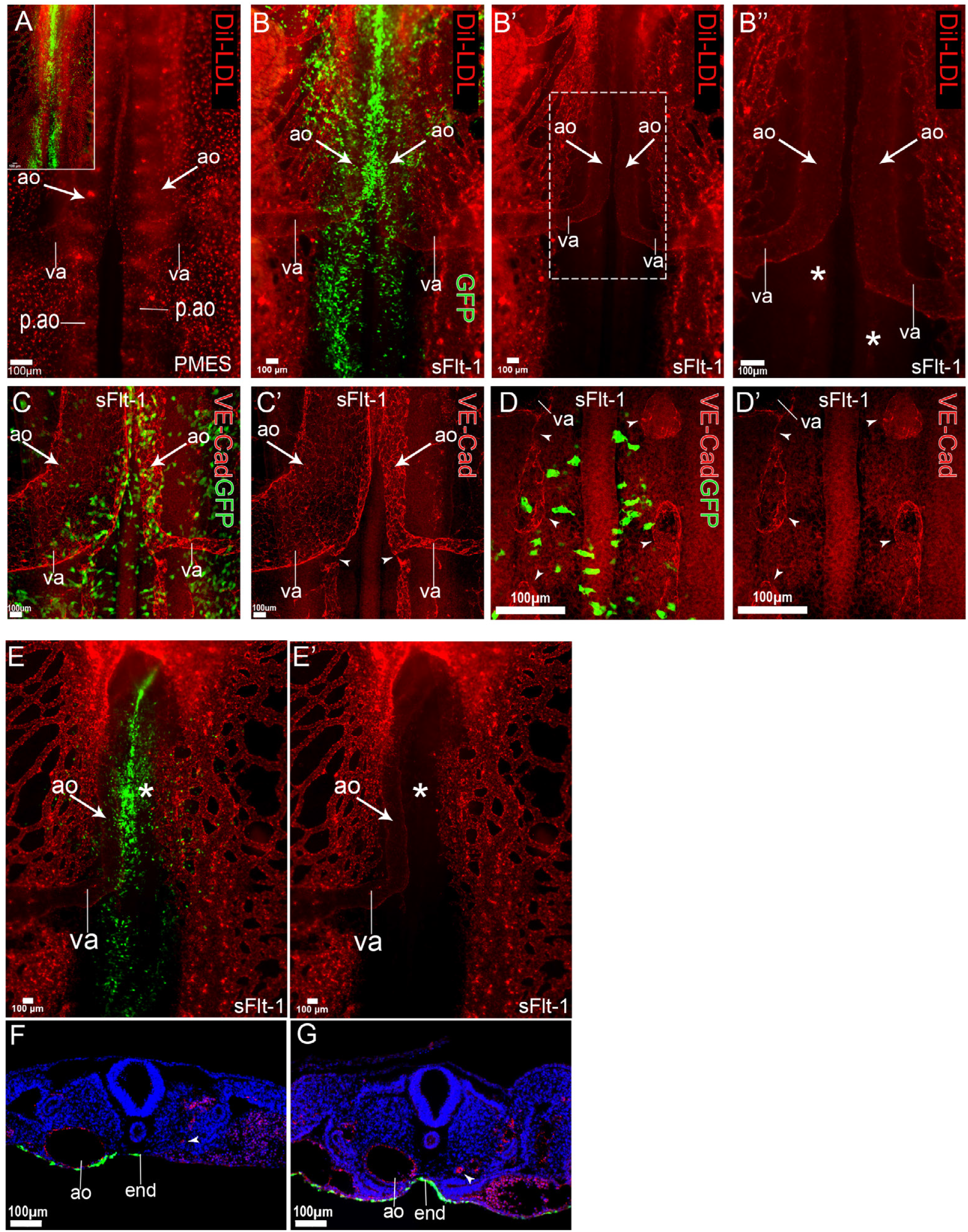
Additional effects of lowering VEGF concentrations on aorta morphogenesis. (A-D) Embryos electroporated with control pMES (A) or sFlt1 (B-D) plasmids and imaged in whole mount for Di-LDL (A,B,B’,B’’), VE-Cad (C,C’,D,D’) and Gfp (A inset,B,C,D). Note lack of continuation of a patent aorta posterior to the vitelline artery (* in B’’). In C,C’ note the presence of endothelial cell cords posterior to the va, but these are not connected to the aneterior aorta (arrowheads). (E-G) Embryos electroporated with sFlt1 plasmid and imaged in whole mount for DiI-LDL (E,E’) and Gfp (E), or in section after staining for VE-Cad, Gfp, and Dapi (F,G). Note absence (E’ asterisk, F arrowhead) or severe reduction (G arrowhead) of right aorta. Ao, aorta; end, endoderm; p.ao, aorta posterior to 22^nd^ somite; va, vitelline artery.

This points to the junction of the aorta with the VA as a critical point that is sensitive to levels of VEGF signaling. Another observation was that often sFlt1 electroporation caused asymmetric interference with aorta morphogenesis, and that when this occurs it is consistently the left aorta that is affected (Fig. 4C,E), even though sFlt1 is expressed symmetrically. In those cases when the left aorta is absent (which appears on the right side in the figure), there is typically a meshwork of endothelial cells in its place (Fig. 4C), reminiscent of an earlier phase of aorta formation.

In order to begin to gain insight into the cellular/molecular mechanisms of the effect of VEGF on aorta morphogenesis, we examined aortic endothelial cells using high resolution confocal microscopy. We observed that in aorta of control embryos, the endothelial-specific cell-cell adhesion protein VE-Cad was localized predominantly at cell-cell junctions (Fig. 5A,B). In contrast, in VEGF-electroporated embryos, significant VE-Cad was found in the cytoplasm in addition to cell-cell junctions (Fig. 5C,D). Electroporation of sFlt1 did not affect intracellular VE-Cad distribution, producing similar effects as electroporation of control plasmid (Fig. 5E,F).

**Figure 5.**
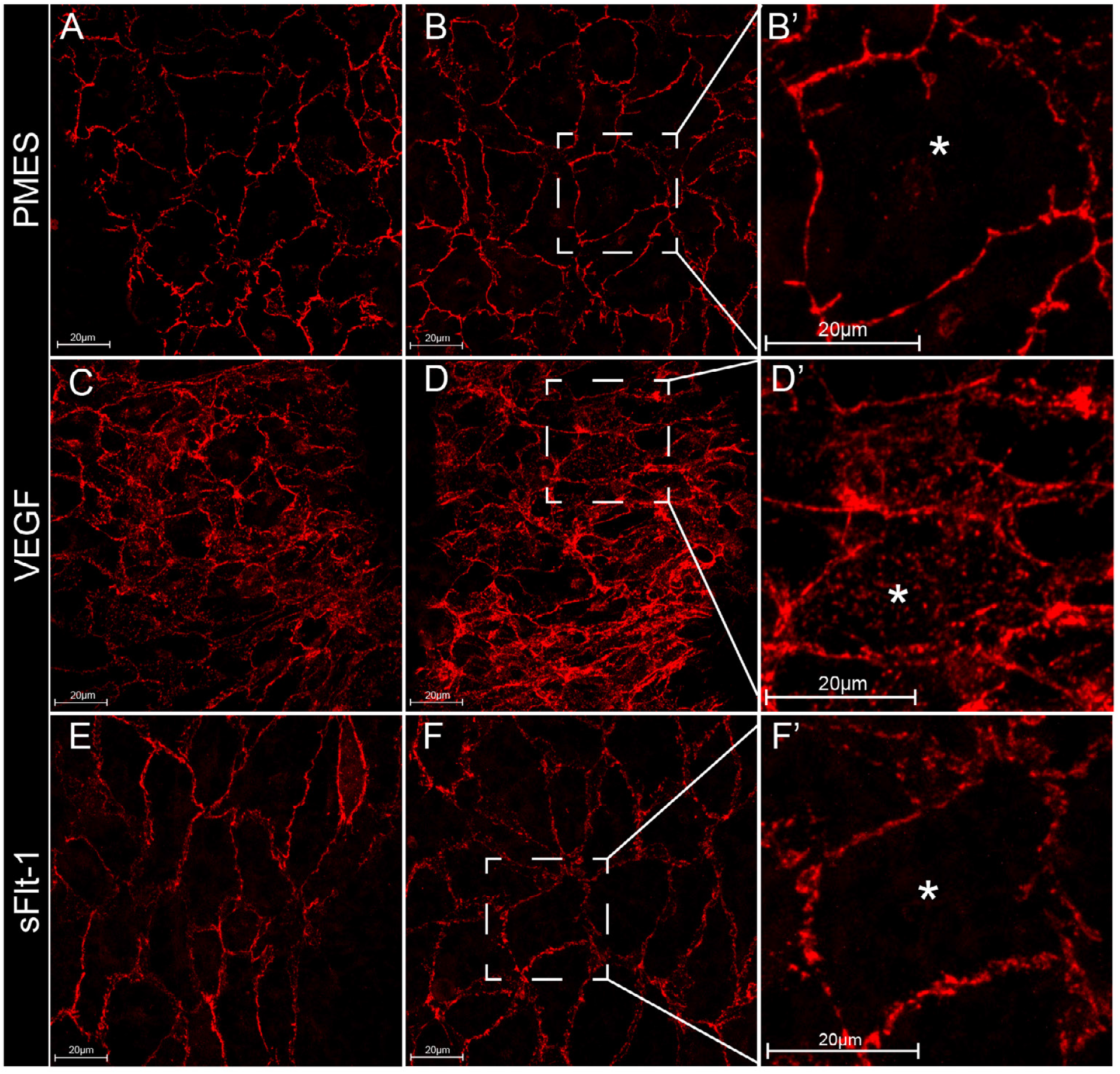
VEGF causes cytoplasmic accumulation of VE-Cadherin. High powered confocal views of the ventral wall of aortas stained for VE-Cadherin after electroporation into the endoderm of control plasmid PMES (A,B), VEGF (C,D), or sFlt1(E,F). Note the accumulation of cytoplasmic of VE-Cadherin in the VEGF-electroporated embryos.

We considered two possibilities for the effect of VEGF on VE-Cadherin. VEGF could be raising VE-Cad expression levels and overwhelming the cellular mechanisms that localize VE-Cad to the cell membrane, with the result that it accumulates in the cytoplasm.

Alternatively, VEGF could be causing a redistribution of VE-Cad from the membrane to the cytoplasm. In order to distinguish between these possibilities, VEGF- or control-electroporated embryos were dissociated into single cells, stained with anti-VE-Cadherin antibodies, and analyzed using Flow Cytometry. We found that, while VEGF produced a significant increase in the number of VE-Cad-expressing cells per embryo, the levels of VE-Cad per cell in VEGF vs. control embryos was the same (Fig. 6). This suggests that VEGF induces a relocalization of VE-Cad away from cell-cell junctions.

**Figure 6.**
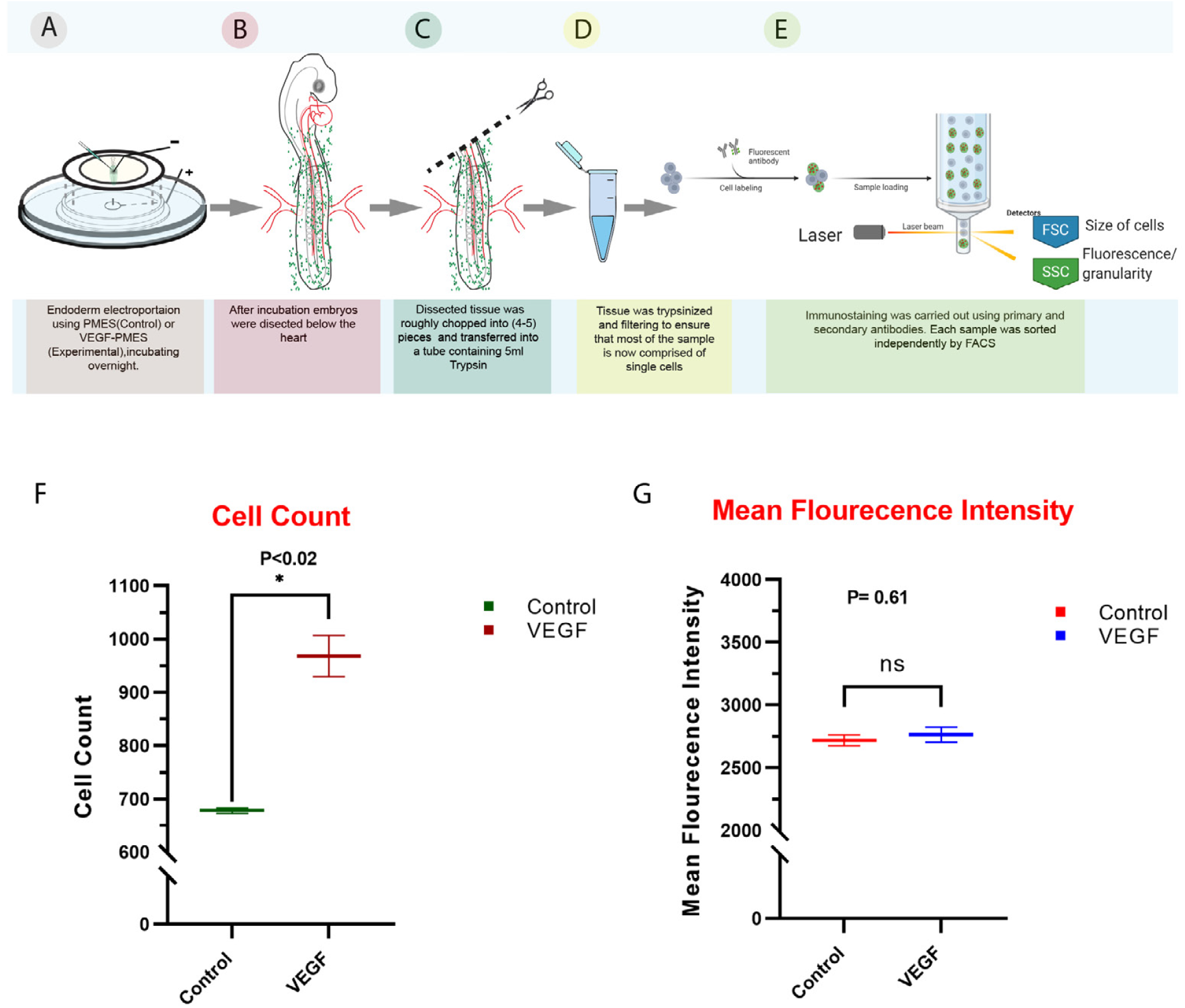
VEGF does not elevate cellular levels of VE-Cadherin. A-E: Experimental procedure. After electroporating the endoderm with VEGF-expressing or control plasmid, embryos were dissociated into single cells, labeled with anti-VE-Cad antibody, and analyzed by Flow Cytometry. F,G: Flow Cytometry Analysis. VEGF caused an increase in number of endothelial cells per embryo (F), but the level of VE-Cad per cell did not change. Bars indicate standard error of the mean.

## Discussion

The current study finds that VEGF levels are critical for proper morphogenesis of the embryonic aorta. Under conditions of lowered VEGF signaling, the aortae degenerate, fail to grow, and fail to migrate towards the embryonic midline. These results are consistent with previous findings indicating that VEGF signaling is required for embryonic blood vessel formation (Ferrara et al., 1996; Fong et al., 1995; Shalaby et al., 1995). In contrast, and consistent with previous studies (Drake and Little, 1995), under conditions of increased VEGF signaling the aortae grow excessively and disproportionally to the rest of the embryo. This growth is caused at least in part by an increase in the number of aortic endothelial cells (Fig. 6E), but an increase in endothelial cell size may also contribute to aortic growth.

Previous studies in Xenopus embryos have uncovered a VEGF requirement for medial migration of bilateral aortic precursors in order to form the single aortic vessel located at the embryonic midline (Cleaver and Krieg, 1998). In Xenopus, VEGF is secreted from the hypochord, a transient structure located at the embryonic midline ventral to the notochord (Cleaver and Krieg, 1998), and VEGF secretion from the hypochord is proposed to act as a chemoattractant that induces migration of aortic cells towards the midline. Amniote vertebrates do not possess a structure resembling a hypochord. The current results find that VEGF is expressed broadly in the area surrounding the aorta, including both medial and lateral to the bilateral aortae, and thus it is difficult to understand how it could act as a midline attractant in the avian embryo. One possibility is that VEGF promotes motility of endothelial cells in a general manner. Since medial regions of the embryo, particularly surrounding the notochord, are less crowded with cells, increased motility could promote migration into less dense regions, as has been proposed in other contexts, such as elongation of the trunk (Benazeraf et al., 2010).

The current results find that VEGF influences intracellular localization of the endothelial cell-specific adhesion molecules VE-Cadherin. Under conditions of increased VEGF, the total amount of VE-Cadherin per cell does not change, but there is a decrease in the proportion of VE-Cadherin that is membrane-localized and a corresponding increase in the presence of VE-Cadherin in intracellular locations. Previous research in cultured cells has reported that VEGF signaling leads to endocytosis of VE-Cadherin (Gavard and Gutkind, 2006; Hebda et al., 2013). Since VE-Cadherin has been found to repress signaling through VEGF receptors (Carmeliet et al., 1999), removal of VE-Cadherin from the membrane could promote signaling through the VEGF pathway. The current results are consistent with this mechanism and suggest that it may be operative in an in vivo embryonic context. Removal of VE-Cadherin from the cell membrane has been found to increase endothelial cell permeability (Corada et al., 1999; Crosby et al., 2005; Gavard, 2014). In the current context, lowering of membrane VE-Cadherin levels may reduce cell-cell adhesion, thereby promoting cell motility and contributing to the remodeling that occurs during aorta migration and fusion.

The effects of effect of lowering VEGF were not uniform amongst the embryonic blood vessels. Some vessels showed marked degeneration or failure to develop, whereas others were relatively unaffected despite high levels of the electroporated VEGF antagonist sFlt1 in the immediate surroundings. With respect to the aorta, the left aorta was consistently more affected than the right, to the extent that in some cases the left aorta was totally absent while the right aorta was present. While the reason for this differential sensitivity is not currently clear, it is interesting to consider that the left and right aortae may experience different hemodynamics and wall stresses due to their positions relative to the outflow tract of the heart (Culver and Dickinson, 2010; Hove et al., 2003; Midgett et al., 2015). Blood flow and wall stress are important factors in vascular remodeling, and that low flow states can promote vessel degeneration (Campinho et al., 2020; Duchemin et al., 2019; Pries and Secomb, 2014). Lowering VEGF signaling may lower aortic growth and blood flow sufficiently that it leads to vessel degeneration on one side, while on the other side the stronger flow rates are sufficient to maintain vessel integrity. Indeed, once the aorta on one side begins to degenerate, this would increase flow rates in the other aorta, thereby maintaining the vessel. Similarly, we observed that the junction between the aorta and the vitelline artery that leads to the yolk sac is highly sensitive to VEGF levels (Fig. 4). In most Sflt1-treated embryos, the connection between the anterior and the posterior aorta is lost, while the connection between the aorta and the vitelline artery is maintained. This suggests that this junction is particularly sensitive to factors that influence vessel growth and maintenance. At this stage of embryogenesis, the posterior regions of the embryo are not yet fully developed and receive relatively low rates of blood flow, whereas the extraembryonic regions are more developed and the vitelline arteries exhibit high flow rates. Reducing VEGF signaling via Sflt1 electroporation may lower flow rates in the posterior aorta sufficiently to lead to degeneration, while in the vitelline arteries flow remains sufficient to maintain vessel integrity. In addition to flow, there may be intrinsic differences between blood vessels with respect to their sensitivity to VEGF levels, as has been suggested from reports of genetic manipulation of VEGF signaling (Li and Ferrara, 2022).

## Methods

### Expression plasmids

pMES drives constitutive expression of an inserted gene and co-expresses eGFP under an IRES element(Swartz et al., 2001). pMES-2 was constructed by addition of a group of unique enzyme sites (NotI, RsrII, AgeI, EcoRV, Eco53kI, SacI, XhoI, AflII, MluI) into the polylinker site into the original pMES plasmid. pMES-Sflt-1 was constructed by subcloning the full human Sflt-1 from NSPI-Sflt-1 (generously provided by G Neufield) into pMES. pMES 2-hVEGF165 was constructed by subcloning human VEGF165 from the pCDNA3A-hVEGF165 plasmid (a generous gift from G. Neufeld) into pMES-2.

### DiI-Ac-LDL

Acylated low density lipoprotein labeled with the fluorescent probe DiI (Biomedical Technologies) was injected directly to the beating heart of the embryo in order to label embryonic blood vessels.

### Electroporation

HH Stage 11-12 embryos were electroporated in an electroporation chamber (James and Schultheiss, 2005) using a 2mm square cathode electrode (NapaGene). Embryos were arranged endoderm up in the chamber and 0.3-2mg/ml plasmid in 5% sucrose and 0.1% Fast Green was deposited onto the endoderm in the following concentrations: pMES= 2000ng; pMES-sFlt1=1600ng; PMES2-VEGF=300-600ng; Rescue=(1600ng pMES-sFlt1 +300-600ng PMES2-VEGF). Three pulses of 12V, 10msec, interval 1sec between pulses were delivered using a BTX830 Square Wave Electroporator (Harvard Biosciences). Embryos were incubated overnight until Stage 16-17 and then processed for immunofluorescence or DiI-LDL injection.

### Immunofluorescence and Clearing

Embryos were fixed in 4% Paraformaldehyde in PBS overnight at 4 degrees. For whole mount immunofluorescence, embryos were permeabilized the embryos with 0.5% Triton X-100 in PBS followed by incubation with primary antibody in TXBuffer (0.2% gelatin, 300mM NaCl, 0.3% triton X-100 in PBS) for two days, washing with 0.3% triton X-100 in PBS and secondary antibody incubation in TXBuffer for two days. Dapi (Sigma) was added at 1mg/ml to the secondary antibody to mark nuclei. Following secondary antibody incubation, embryos were washed with 0.3% triton X-100 in PBS and then three washes are done with PBS. The following primary antibodies were used: mouse anti-GFP (1:500, Molecular Probes 3E6) and rabbit VE-Cadherin (1:500)(Arraf et al., 2017). Secondary antibodies used were Alexa488 and Cy3(1:250, Jackson ImmunoResearch). Some embryos were embedded in 15% Sucrose/7.5% Gelatin/PBS, and cryosectioned at 10um thickness, mounted using Fluorescence Mounting Medium (DAKO S302380), and photographed with either a Zeiss AxioimagerM1 widefield epifluorescence microscope and a Qimaging ExiBlue monochrome camera, or with a Panoramic 250 Flash III Slide scanner (Histech) 250 Flash III. Other embryos were cleared with RapiClear RI=1.47 (SUNJIN Lab) for 1 hour, mounted on a glass coverslip in a chamber with walls constructed from two-side tape, and imaged using a Zeiss LSM880 confocal microscope and 25x and 63x long working distance PlanApo objectives (WD=0.5mm). Image overlays were performed with ImagePro Plus software (Mediacy) or ImageJ.

### In Situ Hybridization

Digoxigenin whole mount in situ hybridization was carried out essentially as previously described (James and Schultheiss, 2005). The VEGFA probe was a 600bp fragment amplified using PCR from HH stage 15 embryo cDNA using primers CCATGAACTTTCTGCTCACTTGG and CTGCTCACCGTC-TCGGTTTTTC and cloned into pGEM-T Easy (Promega). The VEGFR2 probe was a generous gift from C. Kalcheim (Eichmann et al., 1993). Following development of the colorometric reaction, embryos were cryosectioned at 20mm intervals and photographed with a Zeiss AxioimagerM1 widefield microscope with DIC optics and a Qimaging ExiBlue monochrome camera with RGB color filter. For combined immunofluorescence and in situ hybridization, in situ hybridization was performed first, followed by cryosectioning at 20mm intervals and immunofluorescence as described above.

### Flow Cytometry

HH Stage 11-12 chick embryos were electroporated in the endoderm with pMES or pMES-2-vegf. Embryos were incubated for 14 hours (until approximately HH St. 16). After incubation embryos were injected into the beating heart with DiI-LDL, incubated for a further 30 minutes, and then harvested and transferred into culture dishes containing chilled 1xPBS. Embryos were dissected on a silicone plate using a microscalpel (Feather) to cut out the desired area (from just posterior to the heart to the posterior end of the embryo, including both sides of the embryo). The dissected tissue was transferred into chilled DMEM in a small tissue culture plate. Each sample was roughly chopped into 4-5 small pieces and transferred, with minimal DMEM into a 50 ml tube containing 5ml Trypsin (0.25%, Biological Industries), on ice. Trypsin treatment was performed at 37C first in a water bath for 10 minutes and then 20 minutes in a rotary shaker (80 RPM). The reaction was stopped by adding 1ml chilled FBS. A 5ml tip was used to triturate the sample; when no visible tissue chunks were seen the sample was filtered first through a 70mM cell strainer and then through a 40mM strainer into a 50ml tube, and then pelleted in a swinging bucket rotor, at 500G for 7 minutes. The supernatant was carefully aspirated and each sample was resuspended in 750ml of FACS solution (1xPBS, 2% FBS, 2mM EDTA pH8) and held on ice. Samples were then fixed by adding 1ml of 1%PFA in PBS for 15 minutes, followed by centrifugation for 5 minutes. The pelleted cells were washed with cell staining buffer (0.5%BSA in PBS), centrifuged, and cells were permeabilized by adding 2ml permeabilization buffer (0.1% saponin in cell staining buffer) and centrifuged at 600g for 5 minutes at RT. The permeabilized cells were stained with mouse primary antibodies rabbit anti-Ve-cadherin (1:500) and mouse anti-myosin II (DSHB, 1:50) diluted in permeabilization buffer for 20 minuets at 4C. Cells were then pelleted, washed with permeabilization buffer and stained with secondary antibody (Dylight488 Donkey anti-rabbit IgG and AlexaFluor647 Donkey anti-mouse IgG diluted in permeabilization buffer for 20 minutes at 4 degrees C in the dark. Stained cells were washed with permeabilization buffer three times and resuspended in 250ul cell staining buffer. For flow cytometry analysis, cells were gated base on FCS and SSC to exclude debris. Within the gated cells, a doublet discrimination gate was done base on FCS-W/FCS-H and expression analysis analysis was performed on single cells.

### Computer software

Graphs in for Fig. 6 were generated with Prism software (Graphpad). Diagrams were produced using Illustrator (Adobe) and Biorender software.

## Acknowledgements

We thank Gera Neufeld for providing sFlt1 plasmid, and Maya Holdengraber and Melia Guruwitz from the Interdepartmental Imaging Facility for assistance with imaging. This work was supported by grants to T.M.S. from the Israel Cancer Research Fund, the Israel Science Foundation (1463/16), and the Rappaport Family Foundation.

## Data Availability

The datasets generated during and/or analysed during the current study are available from the corresponding author on reasonable request.

